# Over the rainbow: a practical guide for fluorescent protein selection in plant FRET experiments

**DOI:** 10.1101/734616

**Authors:** Grégoire Denay, Patrick Schultz, Sebastian Hänsch, Stefanie Weidtkamp-Peters, Rüdiger Simon

## Abstract

Receptor-like kinases (RLK) and receptor-like proteins (RLP) often interact in a combinatorial manner depending on tissue identity, membrane domains, or endo- and exogenous cues, and the same RLKs or RLPs can generate different signaling outputs depending on the composition of the receptor complexes they are involved in. Investigation of their interaction partners in a spatial and dynamic way is therefore of prime interest to understand their functions. This is however limited by the technical complexity of assessing it in endogenous conditions. A solution to close this gap is to determine protein interaction directly in the relevant tissues at endogenous expression levels using Förster resonance energy transfer (FRET). The ideal fluorophore pair for FRET must, however, fulfil specific requirements: (i) the emission and excitation spectra of the donor and acceptor, respectively, must overlap; (ii) they should not interfere with proper folding, activity, or localization of the fusion proteins; (iii) they should be sufficiently photostable in plant cells. Furthermore, the donor must yield sufficient photon counts at near-endogenous protein expression levels. Although many fluorescent proteins were reported to be suitable for FRET experiments, only a handful were already described for applications in plants. Herein, we compare a range of fluorophores, assess their usability to study RLK interactions by FRET-based fluorescence lifetime imaging (FLIM) and explore their differences in FRET efficiency. Our analysis will help to select the optimal fluorophore pair for diverse FRET applications.

**One-sentence summary:** We compared the performances of several different fluorescent protein pairs to study membrane protein interaction in plants with FRET.

## Introduction

Receptor-like kinases (RLK) are essential components of the plant signaling machinery. They serve to coordinate developmental processes, pathogen recognition, symbiotic interaction with beneficial microorganisms, or other aspects of environmental sensing (Osakabe et al., 2013). RLKs usually act in complexes consisting of a receptor and a co-receptor, and can form higher-order complexes that rearrange dynamically upon ligand perception (Wan et al., 2019). The same receptors can interact with different partners and be part of different complexes with different signaling specificity (Liebrand et al., 2014; Bücherl et al., 2017; Wan et al., 2019). Most studies of RLK interaction initially relied on *in vitro* experiments, and could not provide insights into the spatial localization and complexity of interactions. However, advances in imaging techniques now allow to study co-localization and interactions of RLKs in living plant cells (Somssich et al., 2015; Bücherl et al., 2017). In particular, Förster Resonance Energy Transfer (FRET) is an attractive technique as it allows to resolve protein-protein interactions in live plants in a cell-specific and dynamic manner (Somssich et al., 2015; Long et al., 2017; Weidtkamp-Peters and Stahl, 2017; Lampugnani et al., 2018).

Determination of protein interaction by FRET relies on the energy transfer from a fluorescent donor to an acceptor fluorophore. This phenomenon leads to fluorescence quenching of the donor and excitation of the acceptor, which can be relaxed by fluorescence emission (Förster, 1946). FRET requires that several conditions are met: (i) The emission spectrum of the donor must overlap with the excitation spectrum of the acceptor, (ii) both molecules must be in close proximity (typically <10nm) to each other, and (iii) the dipole moments of both fluorophores must be parallel (Förster, 1946; Clegg, 2006; Bajar et al., 2016b). FRET can be measured in several different ways including intensity-based methods, spectral recording, photobleaching, anisotropy, or fluorescence lifetime (Pietraszewska-Bogiel and Gadella, 2011; Bajar et al., 2016b). Each molecular complex exhibits a specific FRET efficiency, which depends in part on relative fluorophore distance and orientation. However, as FRET is measured through diffraction-limited microscopy, only the apparent FRET efficiency, which represents a mean FRET efficiency in the observation volume, is experimentally accessible (Bajar et al., 2016b). We will refer to the apparent FRET efficiency in the following as “FRET efficiency”, by extension.

Fluorophore pairs for FRET experiments should have a considerable overlap between their emission and excitation spectra, yield a sufficient brightness to fit the fluorescence decay and limit background fluorescence, allow proper folding of the fused protein, do not affect localization, and do not trigger artefactual interactions. There is now a large choice of available fluorescent proteins (Lambert, 2019), however, most of these fluorophores were characterized solely *in vitro* or in mammalian cell systems (van der Krogt et al., 2008; Bajar et al., 2016b). The plant community focused mostly on the use of the GFP-RFP pair (and their derivatives eGFP and mCherry) for FRET experiments (Lampugnani et al., 2018). Additionally, CFP-YFP and other pairs based on improvements upon CFP or YFP were used in plant systems, notably for the Cameleon calcium sensors (Kanchiswamy et al., 2014). Only one study assessed the quality of different FRET pairs in plants and focused on using SYFP2 either as an acceptor for SCFP3A or mTurquoise, or as a donor for mStrawberry, mCherry, or mRFP (Long et al., 2018). In this study the root expressed transcription factors that were analyzed are relatively tolerant to fusions with additional protein domains. However, many membrane proteins consisting of domains with very different properties appear to be more sensitive.

In order to test the suitability of a set of genetically-encoded fluorophores for FRET-FLIM applications with membrane localized proteins in plants, we chose the CORYNEΔkinase (CRNΔKi) -- a kinase-deleted version of CRN -- and CLAVATA2 (CLV2), a receptor-like protein with extracellular leucine-rich-repeats (LRRs) from Arabidopsis thaliana as a test system. Both are membrane localized due to their transmembrane domains, and their interaction *in vivo* has been reported in a number of previous studies (Bleckmann et al., 2010; Kinoshita et al., 2010; Zhu et al., 2010; Breda et al., 2019). CRN and CLV2 are obligate heteromers, and their interaction is necessary for export form the endoplasmic reticulum (ER) to the plasma membrane (PM) (Bleckmann et al., 2010; Somssich et al., 2016). Fusions of fluorophores such as eGFP and mCherry to the cytoplasmic domains of CRNΔKi and CLV2, respectively, showed a very high FRET efficiency (Bleckmann et al., 2010; Somssich and Simon, 2017). For these reasons, CLV2/CRN form an attractive test system to compare the performances of different FRET pairs to study plant membrane proteins interaction: high FRET efficiency can yield a more sensitive readout for comparison, and inactive fusion proteins unable to interact with their RLK partner will remain localized in the ER. To objectively compare different fluorophore pairs for plant FRET assays of RLK, we observed parameters such as localization, photo-stability, and FRET efficiency for several different FRET pairs, including several so far untested combinations in plant systems.

## Results

### Optimization of Fusion Protein Co-Expression in Nicotiana

A quick, easy, and reliable way to test protein-protein interaction by FRET in plants is to use transient expression systems in *Nicotiana benthamiana* (thereafter Nicotiana). For this, both fluorescently tagged proteins must be co-expressed in the same cells. A classical way of co-expressing proteins in Nicotiana is to infiltrate leaves with different plasmids, each carrying the expression cassette for a single protein (Norkunas et al., 2018). This system however presents several problems: co-expression is highly variable, ranging from only a few cells to almost all cells co-expressing both constructs (Hecker et al., 2015); in addition, the relative expression levels of both constructs are very variable from cell to cell (figure 1D). Since FRET is measured with pixel-wise resolution and not at the single molecule level, apparent FRET will differ according to the relative concentration of donor and acceptor (Fábián et al., 2010) (Figure 1A). It is also important to control expression levels as expression under strong promoters can trigger ER stress, protein aggregation and artefactual interactions (Zuo et al., 2002; Bleckmann et al., 2010). To solve the latter problem, we used an estradiol inducible expression system that was previously shown to allow controlled expression of CRN and CLV2 (Bleckmann et al., 2010).

**Figure 1.**
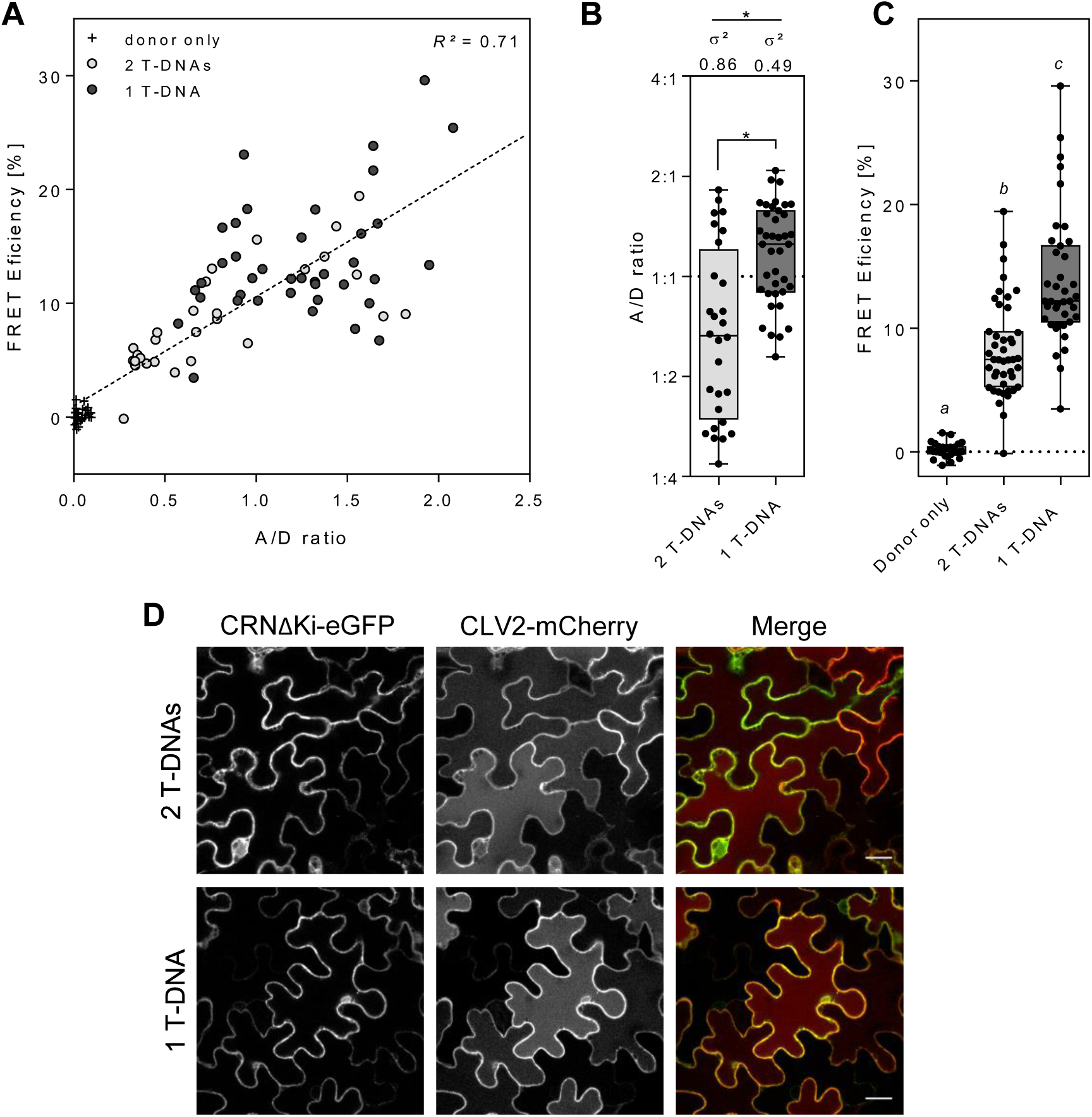
Comparison of the co-expression of fusion proteins from a single or two T-DNAs effect on FRET efficiency. (A) FRET efficiency depends on the expression ratio of acceptor to donor. A/D intensity ratio was calculated as a proxy for the relative level of each fusion protein and plotted against the FRET efficiency of the sample. Both variables are linearly correlated (*R*^2^= 0.71; dotted line). White: expression of CRNΔKi-eGFP only; Dark grey: expression of both CRNΔKi-eGFP and CLV2-mCherry from a single T-DNA; Light grey: expression of CRNΔKi-eGFP and CLV2-mCherry from distinct T-DNAs. (B) A/D ratio (Log2 scale) was significantly higher in samples co-expressing the two fusion proteins from a single T-DNA in comparison to those expressing from two T-DNAs (Tukey, *p*<0.0001, N≥28). The variance (σ^2^) of the two samples was also significantly different (F-test *p*<0.01). (C) FRET efficiency was significantly higher in samples co-expressing the two fusion proteins from a single T-DNA in comparison to those expressing from two T-DNAs (Tukey, *p*<0.0001, N≥29). (D) Co-expression of CRNΔKi-eGFP (Green) and CLV2-mCherry (Red) from a single T-DNA (bottom row) results in a higher coexpression rate than when expressed from 2 T-DNAs (top row). Yellow indicates colocalization of both eGFP and mCherry. Scale bar: 25μm.

With the aim of reducing variability in the ratio of acceptor to donor, we designed an expression vector allowing simultaneous expression of two distinct fusion proteins under the control of the estradiol inducible system from a single T-DNA. We measured both Acceptor/Donor (A/D) fluorescence ratio and FRET efficiency of cells co-expressing CRNΔKi-eGFP and CLV2-mCherry from a single T-DNA and compared them to cells co-expressing from individual T-DNAs (Figure 1B and C). Co-expression from a single T-DNA resulted in a strong reduction in the variance of the A/D ratio (F-test, *p*<0.01; Figure 1B). Surprisingly, the average A/D ratio was higher than when the fusion proteins were co-expressed from independent T-DNAs, although we cannot explain this effect. This resulted in the FRET efficiency being higher as more acceptor was available to quench the donor fluorophore (Figure 1C). Consistent with expectations, nearly all expressing cells co-expressed both constructs when carried on a single T-DNA (Figure 1D). In conclusion, co-expression of both fusion proteins from a single T-DNA proved to be a more suitable system for FRET analysis than the classical way of co-expressing proteins from distinct T-DNAs, for the following reasons: (i) It reduced the variability in co-expression levels, thereby reducing the potential influence of relative protein concentration on apparent FRET and increasing measurement reproducibility. (ii) It increased the apparent FRET. (iii) It greatly improved the co-expression rate, reducing the amount of time spent finding suitable cells. We therefore used the coexpression system for the rest of this study.

### Selection and Expression of Fluorophore Pairs for FRET Measurements

Several fluorophore pairs were previously described to yield efficient FRET in plant or mammalian models (Table 1). We used the above described system allowing inducible transient co-expression of tagged CRNΔKi-donor and CLV2-acceptor fusion proteins from a single T-DNA to assess the quality of these different FRET pairs in Nicotiana cells.

**Table 1.**
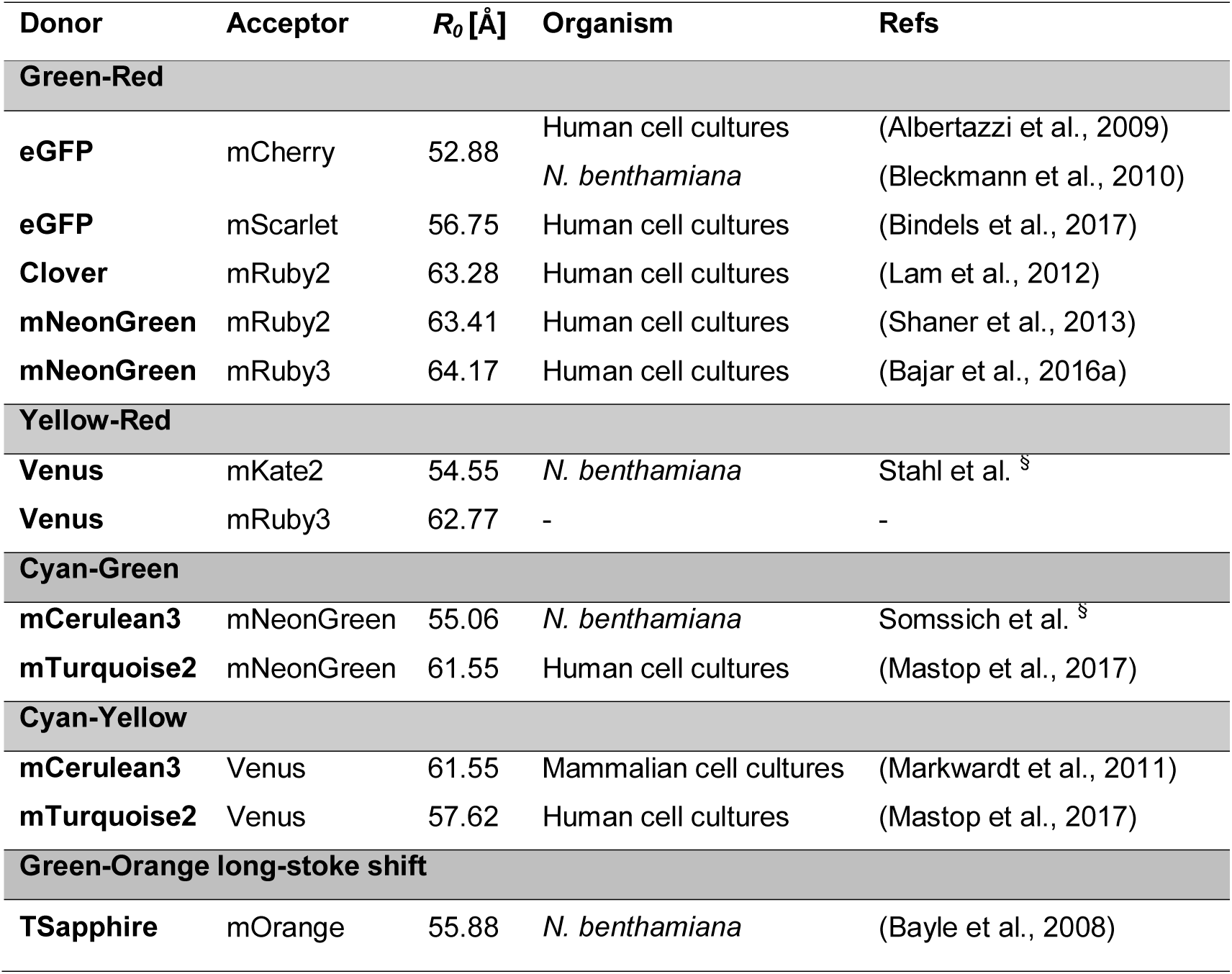
Overview of the FRET pairs used in this study. References for previously published FRET experiments and study organisms used are indicated. R_0_: Förster radius of the FRET pair. §: personal communication.

**Table 2.**
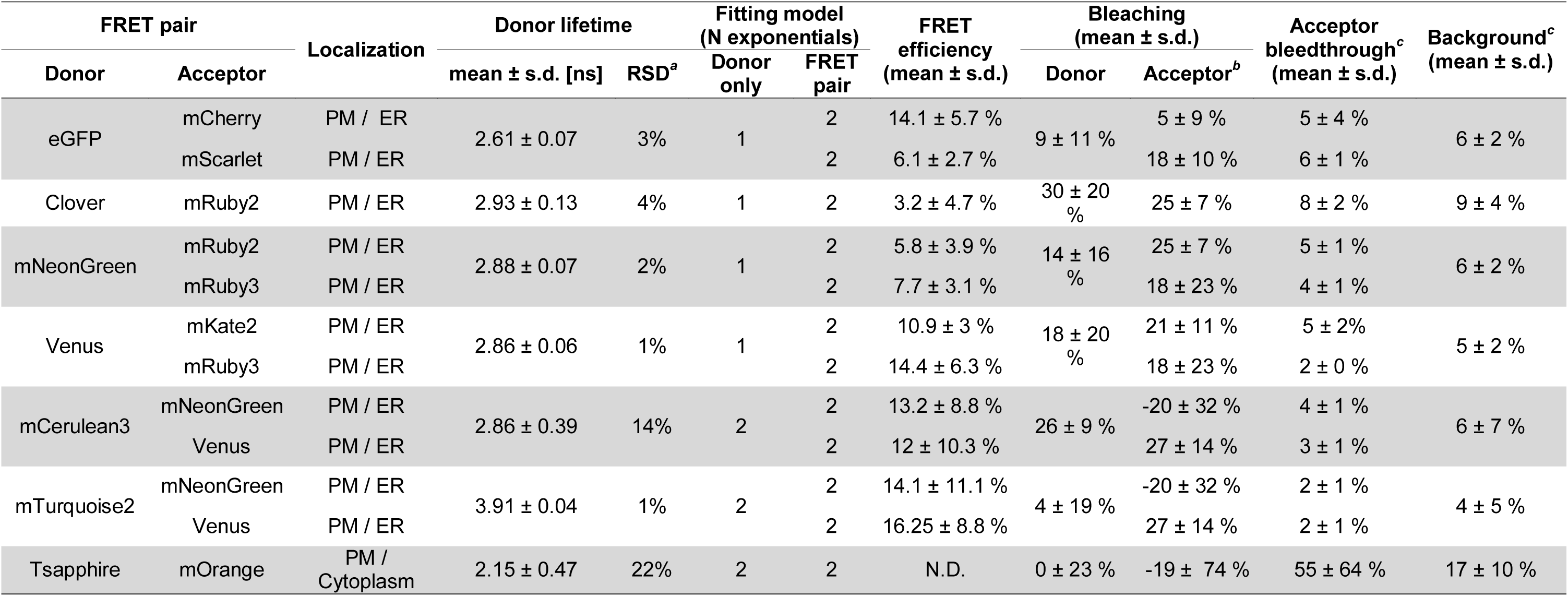
Summary of the FRET pairs examined. a: Relative standard deviation; b: Negative values indicate an increase of fluorescence; c: Given in percent of the mean donor intensity.

FRET requires sufficient rotational freedom of the fluorophores, so that dipole moments can align and permit resonance energy transfer. The conformational state of the fusion proteins is, however, mostly unpredictable. Addition of linker sequences between the fusion protein and the fluorophore can increase the likelihood of FRET (Stryer and Haugland, 1967; Lissandron et al., 2005; Osad’ko, 2015), if the linker increases free rotation of the fluorophore while being short enough to keep the fluorophores in close proximity in case of complex formation. Whether a linker sequence improves FRET efficiency can be experimentally tested for individual protein combinations.

### Fluorophore Effect on Protein Localization

CLV2 and CRN are exported from the ER to the PM only if they interact with each other via their transmembrane domains (Bleckmann et al., 2010; Somssich et al., 2016), and they therefore constitute a convenient system to investigate the correct folding and interaction of fusion proteins.

High levels of protein expression can result in overloading of the vesicular transport system and lead to protein retention in the ER and formation of organized smooth endoplasmic reticulum (OSER) through weak protein-protein interactions (Snapp et al., 2003; Ferrero et al., 2015). For these reasons we used the above-mentioned estradiol-inducible system, allowing controlled expression of the transgene, so that the optimal timing for measurement can be tested for each fusion proteins. It was previously shown that a short induction time of 4h led to predominantly PM-localization of CLV2 and CRN, while longer induction times of more than 12h led to the formation of protein aggregates and predominant ER localization (Bleckmann et al., 2010). However, in our hands overnight expression of CRNΔKi-eGFP and CLV2-mCherry resulted in predominantly PM localization occasionally associated to ER localization, while aggregation was observed in only a few cells (Figure 1D). As FLIM experiments typically require extended amount of time, we used overnight induction and acquired fluorescence lifetime over the course of a full day. Expression of all gene constructs resulted in a minor amount of ER localization of the fusion proteins, in addition to their expected PM localization, but aggregation was rarely observed (Figure 2), indicating that none of the fluorophores significantly interfered with CLV2-CRN interaction.

**Figure 2.**
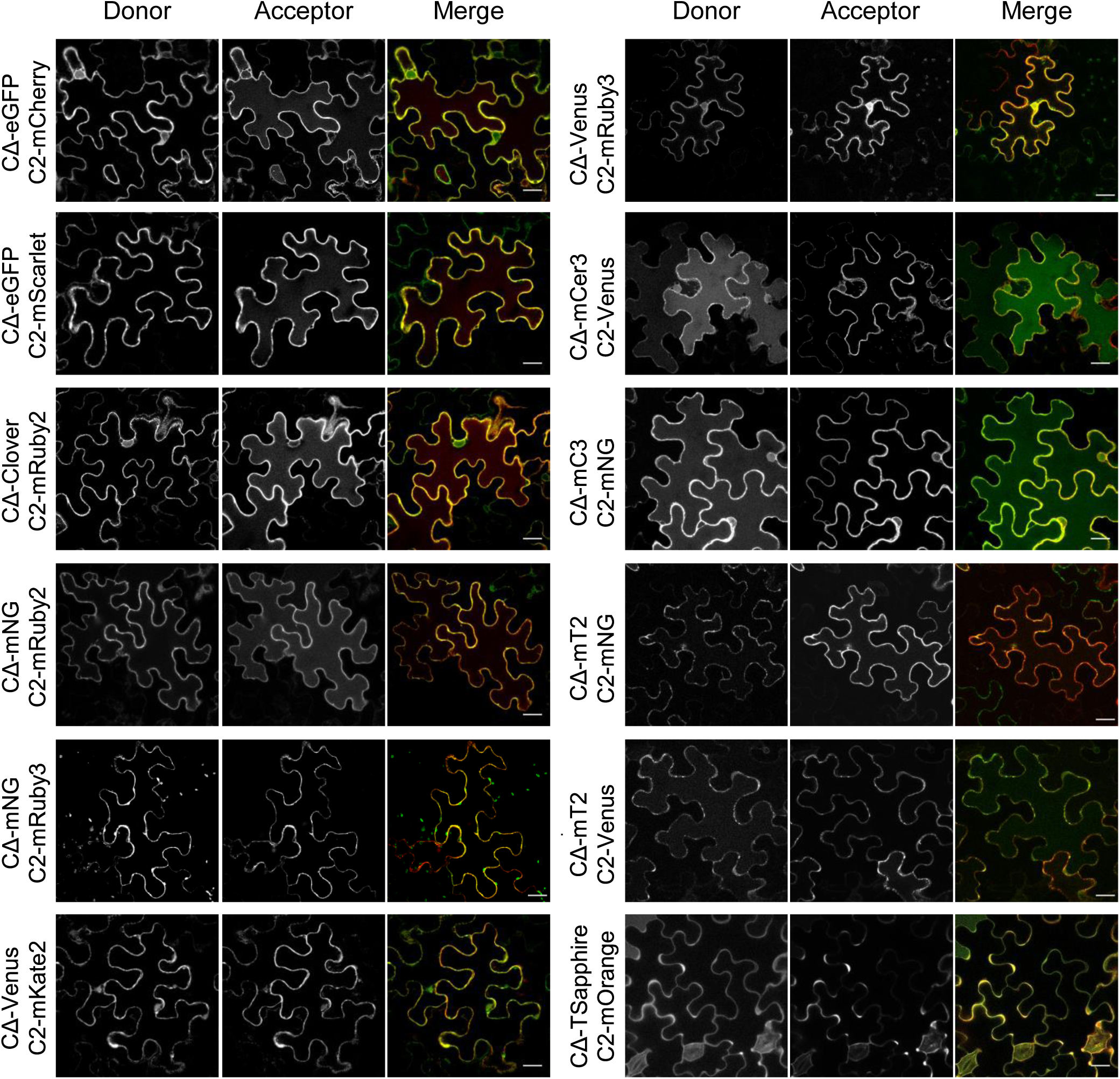
Subcellular localization of the RLK-fluorophore fusion proteins. Confocal microscopy of Benthamiana cells co-expressing CRNΔKi-donor (CΔ) and CLV2-acceptor (C2) fusion proteins. Both donor (left row) and acceptor (middle row) are shown as grey scale and in false colors in the merged images (right row; green: donor; red: acceptor). Scale bar: 25μm. mNG: mNeonGreen; mCer3: mCerulean3; mT2: mTurquoise2.

Fusion proteins with Venus and Clover typically showed higher ER retention than other fluorophores. This may be due to the presence of an Alanine in position 206 of both proteins which is responsible for weak dimerization and the accumulation of membrane-localized fusion proteins in ER structures (Zacharias et al., 2002; Snapp et al., 2003). For this reason, the monomeric variants mClover3 and mVenus should be preferred for membrane proteins fusions. Expression of CRNΔKi-T-Sapphire and CLV2-mOrange were typically weak and showed extensive accumulation in cytoplasmic regions in zones of high membrane curvature. Like Venus and Clover, T-Sapphire contains an Alanine in position 206, enabling it to dimerize. Additionally, T-Sapphire is a fast folding mutant of GFP (Zapata-Hommer and Griesbeck, 2003). Fast folding of the fluorophore might lead to the misfolding of the RLK moiety and accumulation of aggregated, misfolded fusion proteins in specific membrane or cytoplasmic regions.

mCherry, mScarlet, mRuby2, mRuby3, mTurquoise2, and mCerulean3 fusions typically showed a weak fluorescence in the vacuole of expressing cells. This could be a sign of protein recycling or storage in the vacuole and is likely a general property of all fluorophores. As these fluorescent proteins have a low pKa (≤5.3) they are still fluorescent at the slightly acidic vacuolar pH (5.5-6), at which eGFP fluorescence at 488nm is largely quenched (Haupts et al., 1998; Martinière et al., 2013). These less pH-sensitive fluorophores may be useful to study proteins targeted to acidic compartments such as the lytic vacuole or apoplasm.

### Fluorophore Sensitivity to Photobleaching

Energy transfer from the donor fluorophore to the acceptor is impaired by bleaching of the latter. While this is the leveraged mechanism for FRET measurement in acceptor-photobleaching experiments, this can be a disadvantage in FLIM experiments as the measured lifetime will increase in correlation to acceptor bleaching. We therefore quantified acceptor photobleaching during a typical FLIM experiments (Figure 3A), and found mCherry to be the most stable of all red acceptors, with minimal fluorescence intensity loss by the end of the measurement. mScarlet, mRuby3 and mKate2 showed bleaching of about 20% of the initial signal, while mRuby2, and Venus were the most sensitive acceptors with about 25% of signal loss during the course of the acquisition. Surprisingly, mNeonGreen and mOrange appeared to be photoactivated by the 440nm laser used for donor excitation, and their signal intensities increased by over 20% during the measurement. However, while this increase was steady for mNeonGreen, mOrange went through a first bleaching phase before its fluorescence sharply increased.

**Figure 3.**
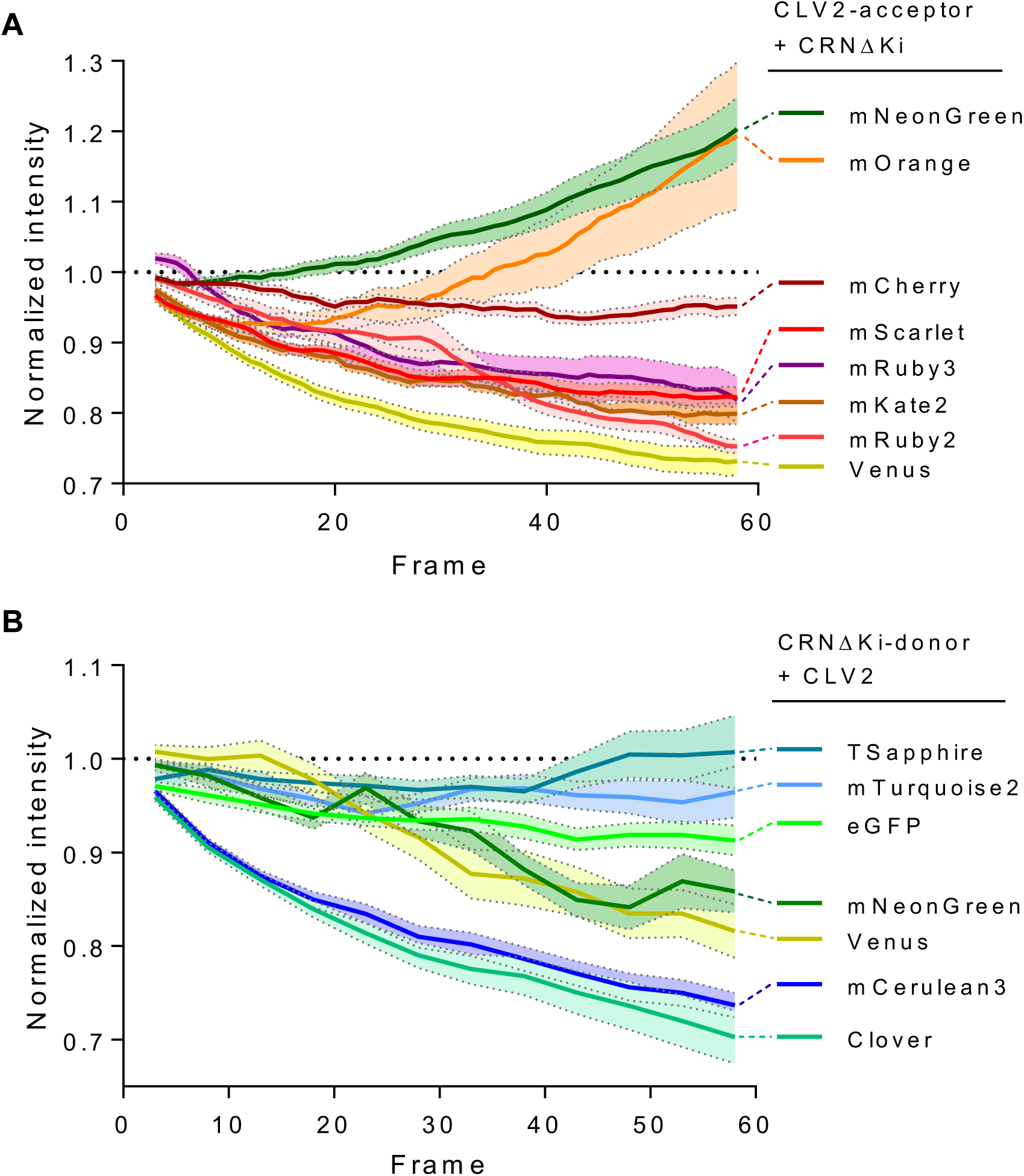
Photobleaching effect during lifetime acquisition. Acceptor (A) and donor (B) photobleaching during lifetime measurement in the absence of their FRET partner. Running average fluorescence intensity of 10 samples were calculated over 5 frames (plain lines). Running standard error of the means are represented as colored areas between dotted lines. Fluorophores are indicated on the right, connected to their final value by a dotted line. Intensity was normalized to that of the first frame.

Donor photo-bleaching during lifetime acquisition decreases the signal-to-noise ratio over time, impacting the quality of the data. Additionally, as bleaching can occur before the relaxation of the fluorophore, and therefore before the emission of a photon, this can artificially decrease the apparent lifetime. We therefore examined donor bleaching over time during lifetime acquisition (Figure 3B). Among the blue-spectral range donors, mTurquoise 2 was very stable (<5% bleaching), while mCerulean3 showed continuous bleaching over the course of acquisition. For the green and yellow donors, TSapphire was the most stable with barely any bleaching over the course of the acquisition. eGFP was relatively stable with less than 10% bleaching, mNeonGreen, and Venus showed mild bleaching of about 15% of the initial fluorescence, while Clover bleached rapidly down to about 40% of the initial fluorescence. Because of their instability, mCerulean3 and Clover should therefore be avoided as donors for FLIM experiments.

### Auto-Fluorescence Background

Plants cells contain numerous compounds such as chlorophyll and phenols that give strong auto-fluorescence in the red and blue range, respectively. As these compounds’ fluorescence lifetime is typically very short, these can lower the apparent lifetime of the sample when their signal is too strong in the donor channels.

We therefore measured fluorescence background of mock-infiltrated Nicotiana for all the different laser and filter combinations that we used (Figure 4). The setups used for the acquisition of Green-Red and Yellow-Red pairs yielded the least background fluorescence, typically one order of magnitude lower than a typical fluorophore measurement. Setups used for the acquisition of the blue-range donors yielded more variable fluorescence which was in some rare cases in the order of magnitude of a typical fluorophore measurement. Autofluorescence originated from the cell wall in these cases and was therefore spatially confounded with the fluorophore signal. Similarly, the setup used for the acquisition of T-Sapphire yielded a relatively high background in comparison to a typical fluorophore measurement, representing around one third of the signal. Nevertheless, auto-fluorescence giving rise to very short lifetime can be filtered out using post-acquisition data filtering (Antonik et al., 2006).

**Figure 4.**
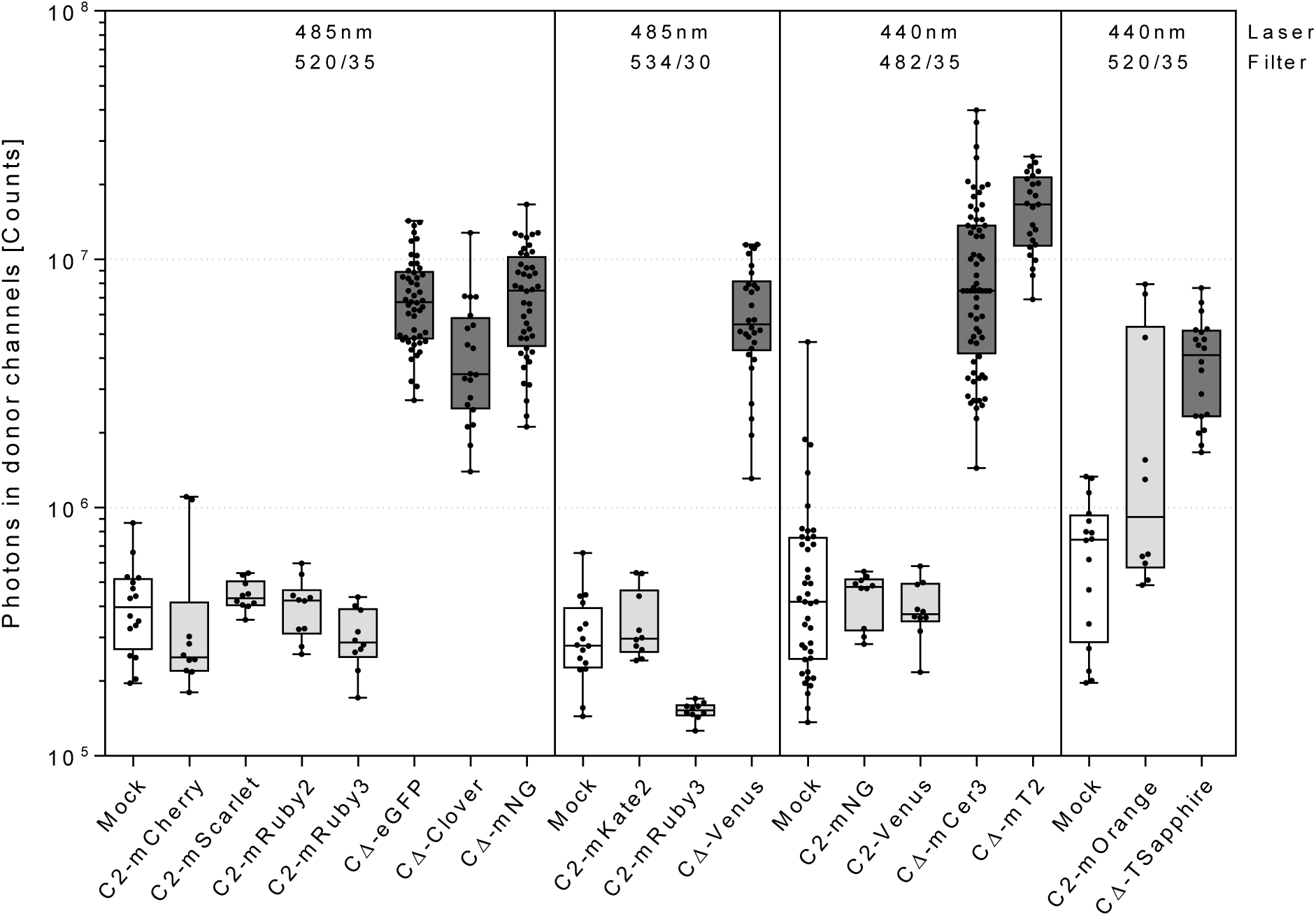
Background and bleed-through controls for the different FRET pairs tested. For each different microscope settings (Laser wavelength and bandpass filter indicated on top of each section), we measured the intensity of the fluorescent background of mock infiltrated plants (white) and the emission of the CLV2-acceptor constructs in the acceptor channels (light grey). Data is represented in comparison to the fluorescence intensity of representative CRNΔKi-donor only constructs (dark grey). Total amount of photons collected are displayed as a log scale.

### Acceptor Bleed-Through Effect

As the signal collected in the donor channels does not discriminate between donor and acceptor emitted photons, it is crucial to determine whether the acceptor-emitted light is able to reach the donor’s channel detectors, albeit the presence of a wavelength-specific bandpass filter. We therefore expressed untagged CRNΔKi together with acceptor-tagged CLV2 and measured the number of photons collected in the corresponding donor channels (Figure 4). In almost all cases the photon counts were similar to that of the background (mock infiltrated Nicotiana). However, mOrange yielded a large photon count in the T-Sapphire channels when excited with the 440nm pulsed laser. As we used a 520/35 bandpass filter before the single photon detectors of the donor channels, mOrange signal should have been intercepted. Although a green-range fluorescence state was never described for mOrange, it is possible that when excited with indigo light, mOrange fluoresces in a green state as described for DsRed, from which mOrange derives (Baird et al., 2000).

### Comparison of Fluorophore Pairs for FRET Efficiency

We next compared the FRET efficiency of the different fluorophore pairs. Therefore, fluorescence decay of the donor was quantified in Nicotiana epidermis cells transiently expressing either CRNΔKi-donor and CLV2-acceptor or CRNΔKi-donor and untagged CLV2 (donor only) with picosecond resolution. We then eliminated background and ER signals by setting a photon minimum count threshold and manually excluding ER and chloroplast-containing regions in order to enrich our samples in PM-localized fluorescence. Fluorescence decays can be well-described by multi-exponential models which allow to extract fluorescence lifetime (τ) as a model’s parameter (Berezin and Achilefu, 2010). As multi-exponential models yield several lifetime components, these need to be weighted and averaged to extract a single τ per sample. For this, each decay component’s τ is weighted by its contribution to the decay’s amplitude. Most donor-only samples could be fitted with a single exponential model to the exception of mCerulean3 and T-Sapphire which required a bi- and tri-exponential model, respectively. Apart from the T-Sapphire-mOrange FRET pair, which required a tri-exponential model, all FRET pair decays could be fitted with a bi-exponential model. FRET efficiency can then be determined as a measure of the reduction of the donor’s lifetime in the presence of an acceptor in comparison to donor only samples. FRET efficiency is typically expressed in percent, with 0% representing the absence of lifetime reduction, and 100% representing total quenching of the donor’s fluorescence.

In order to ensure that lifetime reduction was due to interaction between CRNΔKi and CLV2 and not a consequence of the presence of a large concentration of acceptor surrounding the donor fluorophore at the plasmamembrane, we co-expressed CRNΔKi-eGFP, CLV2-untagged, and a myristoylated version of mCherry (myr-mCherry) anchored in the PM (Turnbull and Hemsley, 2017). CRNΔKi-eGFP and myr-mCherry largely co-localized (Supplementary figure 2). Although the ratio of eGFP to mCherry signal was in the same order of magnitude for the myr-mCherry construct as for the previously described CLV2-mCherry constructs (Supplementary figure 1 and Figure 1), eGFP fluorescence lifetime was not significantly reduced (Figure 5B), indicating that the mere presence of the acceptor at the membrane is not sufficient to quench the donor. Lifetime reduction is therefore a consequence of protein-protein interaction.

**Figure 5.**
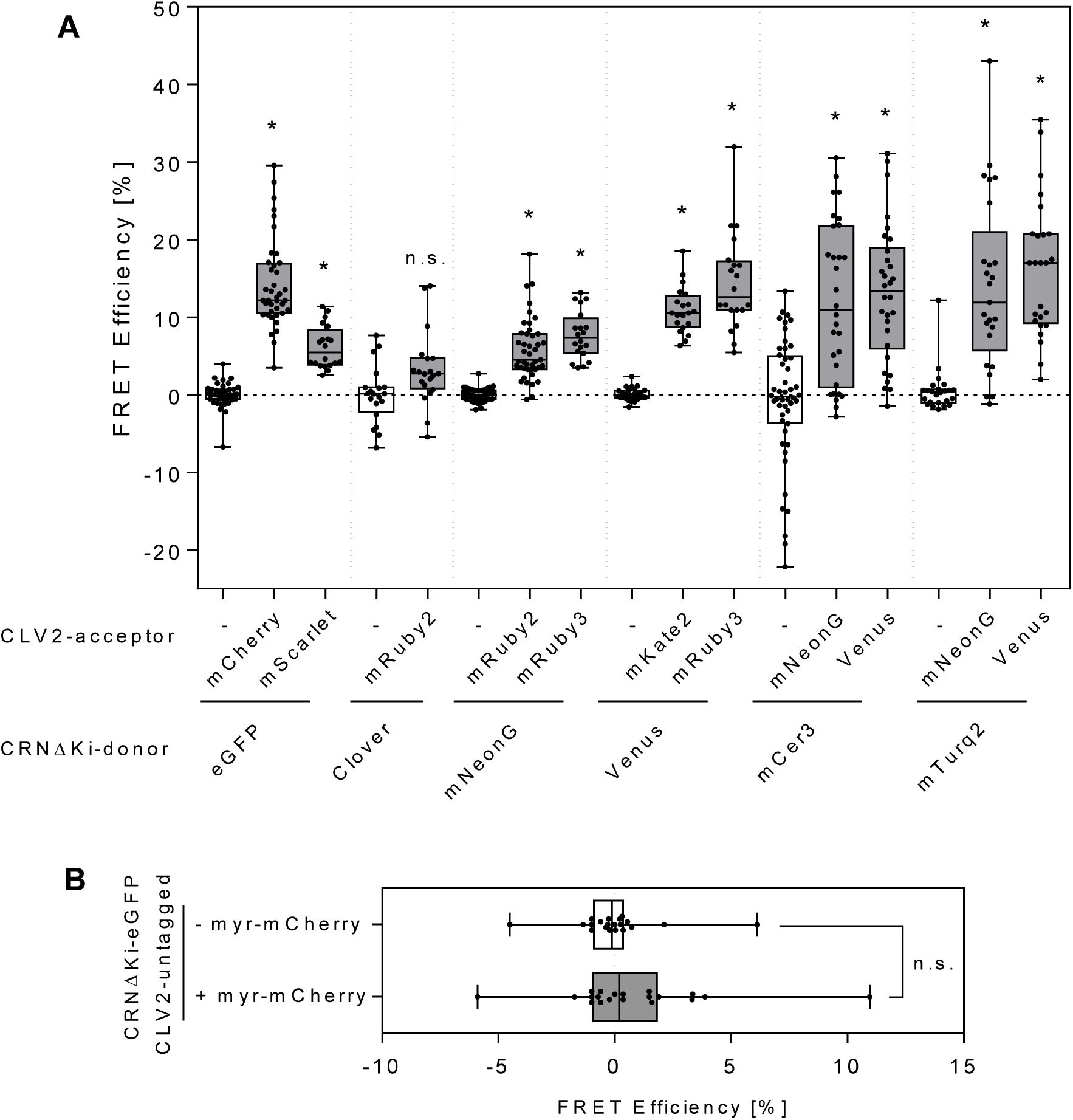
FRET efficiencies of the different FRET pairs tested. FRET efficiencies of the donor only samples (white) and FRET pairs (grey) for CLV2 and CRNΔKi fusion proteins (A) or CRNΔKi-eGFP and plama-membrane localized myr-mCherry control (B). FRET efficiencies were calculated as a normalized measure of the donor’s lifetime reduction. mNeonG: mNeonGreen; mCer3: mCerulean3; mTurq2: mTurquoise2; myr: myristoylation. Stars in A indicate that the sample mean is significantly different to the donor only mean (Holm-Sidak corrected *p*<0.01; N≥20). Absence of difference between the two populations in B was determined by Student T test (*p*>0.1; N=20).

We therefore proceeded to determine the FRET efficiency of our candidate FRET pairs (Figure 5). Surprisingly, we failed to detect any significant FRET between Clover and mRuby2, although this was described as an efficient pair in mammalian cells (Lam et al., 2012). For mCerulean3, we encountered difficulties to consistently fit lifetime (Supplementary figure 2; RSD of 14%), and the average lifetime value we determined (2.9 ns) is inconsistent with the theoretical value of (4.1 ns). Because of the considerable bleed-through of mOrange in the donor channel of TSapphire, the inconsistent localization of both fluorophores, and the high variation of TSapphire lifetime between experiments (Supplementary figure 2; RSD of 22%), we decided to leave the TSapphire-mOrange FRET pair out of our lifetime analysis. They were, however, described as a well performing FRET pair to measure interaction between a transmembrane and a cytosolic protein in another report (Bayle et al., 2008). However, in this report T-Sapphire was fused to a soluble protein, and it is possible that fusion of TSapphire to transmembrane proteins results in mis-folding or -localization. Importantly, the previously described eGFP-mCherry combination was one of the best performing pairs, confirming its status of reference FRET pair for plant experiments. However, combination of Venus with either mKate2 or mRuby3 and using mCerulean3 or mTurquoise2 as donors for either mNeonGreen or Venus yielded comparable FRET efficiencies. The recently described mNeonGreen-mRuby2 and eGFP-mScarlet pair (Shaner et al., 2013; Bindels et al., 2017) yielded measurable FRET, although lower than the eGFP-mCherry pair. mNeonGreen performed significantly better as a donor for mRuby3.

## Discussion

### Improved FRET Measurement from T-DNA Stacking

FRET measurements requires the presence of both donor and acceptor in the same cell, and efficiency is partially dependent on acceptor to donor ratios. As both these factors are highly variable when co-infiltrating donor and acceptor constructs in Nicotiana, we used the GreenGate system (Lampropoulos et al., 2013) to conveniently create a large number of constructs allowing inducible co-expression of donors and acceptors from single T-DNAs, respectively. As previously reported for a similar Gateway-based system (Hecker et al., 2015), this resulted in a drastic increase in co-expression rate as well as a decrease in donor-to-acceptor ratio variability. Additionally, the use of the GreenGate system allows to easily change expression driver (inducible or constitutive, weak or strong), fluorescent tag, and fusion orientation, providing a flexibility unmatched by Gateway based systems. The ability to freely vary these parameters can be leveraged to increase measurements robustness and sensitivity, and time-efficiency of FRET-FLIM experiments.

### Comparison of Fluorophore Suitability in Fusion to RLKs

The optimal fluorophore combination to measure FRET is dependent on the fusion proteins and therefore several combinations could be tried using the present study as a starting point. Fusion protein folding, stability, and activity may be improved by adding amino acid linker sequences of various lengths and composition between the protein of interest and the fluorophore moiety. It is necessary prior for a FRET experiment to verify that the fusion proteins localize properly and are functional.

In addition to the specific protein fusion behaviors, individual fluorophores exhibit different properties, which should be carefully considered before selecting a protein fusion. One point of particular importance for membrane proteins transiting through the ER is to avoid the use of fluorescent proteins that can self-interact, forming oligomers. As membrane proteins are stacked in a single plane, they can reach high local concentration in over-expression situations. In conjunction with weakly self-interacting fluorophores this can lead to the formation of oligomers that convert the ER tubular network into organized smooth ER (Snapp et al., 2003; Costantini et al., 2012). For this reason, monomeric variants such as mEGFP, mVenus, or mClover3 should be preferred. It is also important to confirm that the fusion protein correctly localizes, since some fluorophore fusions can prevent proper folding of their cargo and lead to non-functional and/or mislocalized proteins. For this reason, we decided to exclude the TSapphire-mOrange pair from our lifetime analysis, although they were shown to perform well for soluble proteins (Bayle et al., 2008). Another aspect to be evaluated for individual fluorophores is their signal-to-noise ratio in the cellular context. Imaging of blue-range donors typically yields high cell-wall fluorescence, and very bright fluorophores have to be employed to minimize interference from auto-fluorescence. mCerulean3 and mTurquoise2 typically yielded a very bright signal which counterbalanced the background. It is to note that mTurquoise2 was significantly brighter than mCerulean3 and can therefore be considered a more suitable donor fluorophore for plasmamembrane-localized proteins (Goedhart et al., 2012). Similarly, fluorophores emitting in the far-red spectral range should not be selected for experiments in photosynthetic tissues

Lifetime determination from the fluorescence decay requires a high photon budget, however, as photons are accumulated over several frames weak fluorescence signal can be balanced out by longer acquisition provided that: (i) fluorophores are photostable, (ii) background fluorescence is an order of magnitude lower than the signal, or can be filtered out during downstream analysis (Antonik et al., 2006), (iii) the proteins of interest are slow diffusing and (iv) the tissues of interest do not present significant growth during the measurement.

### Comparison of FRET Pairs Quality

TSapphire and mOrange resulted in mislocalisation of the fusion proteins. Additionally, mOrange emission can affect the detection of TSapphire emission, leading to a bias in TSapphire lifetime determination and should therefore be avoided. Clover and mCerulean3 photobleached significantly during lifetime acquisition; furthermore, we failed to detect FRET between Clover and mRuby2. Therefore, using mCerulean3 and Clover should also be avoided. On the other side we measured FRET between several other pairs. mScarlet did not perform as well as mCherry as an acceptor for eGFP. The eGFP-mCherry pair is preferable when using eGFP as a donor, especially since there is now a considerable amount of literature using this FRET pair. Several good alternatives to this pair exist: (i) Venus can serve as an excellent donor for both mKate2 and mRuby3, where mRuby3 yields a higher FRET efficiency. (ii) mTurquoise2 can be employed as an efficient donor for both mNeonGreen and Venus. mNeonGreen is less sensitive to photobleaching than Venus and will therefore be preferred, although this effect is marginal on FRET efficiency. mNeonGreen gave a reliable FRET as a donor for mRuby3, although yields a lower efficiency than the above-mentioned pairs.

There are several newer and more performant versions of some of the fluorophores we tested here that could further increase FRET performances in plant systems. For example SYFP2 was described to be a brighter variant of Venus which was successfully applied for FLIM measurement of plant transcription factors (Kremers et al., 2006; Long et al., 2018). Likewise, mClover3 is a more stable variant of Clover, that may display better characteristics than its ancestor in plant FRET experiments (Bajar et al., 2016a).

While most FRET applications look at interactions between two partners, a few recent publications pushed this boundary to investigate interactions between three or even four partners within RLK complexes (Breda et al., 2019; Gloeckner et al., 2019). These studies rely on the measurement of competitive interactions or the direct measurement of three-fluorophore FRET-FLIM, both using combinations of three different fluorophores forming overlapping FRET pairs. In the present study we identify two such combinations -- namely mTurquoise2-mNeonGreen-mRuby3 and mTurquoise2-Venus-mRuby3 -- that could further improve these new applications.

## Material and Methods

### Generation of GreenGate Entry Plasmids

Unless otherwise stated, template, destination, and additional entry plasmids used in this work were previously described and are available from the GreenGate cloning kit (Lampropoulos et al., 2013).

The XVE coding sequence was amplified from pABindGFP (Bleckmann et al., 2010) with primers GG_XVE_F and GG_XVE_R (Supplementary table 1), internal *Bsa*I site was removed with PW_XVE_3084_AT_R and PW_XVE_3084_TA_F and the product was cloned in pGGC000 to create pRD71. The LexA-mini35S promoter was amplified from pABindGFP (Bleckmann et al., 2010) with primers GG_EST_F and GG_EST_R and cloned in pGGA000 to create pBLAA001. The coding sequences from CLV2 and CRNΔKi were amplified from pAB125 and pAB128 (Bleckmann et al., 2010) with PS-GG-CDS-Clv2-F and PS-GG-CDS-Clv2-R, and oGD335 and oGD336, respectively and cloned in pGGC000 to create pGD292 and pGD293. The myristoylation sequence (myr) was created by annealing oGD339 and oGD340 before cloning the resulting double-stranded oligonucleotide in pGGC000 to create pGD318. The resistance dummy cassette contains the very short SV40 origin of replication which was amplified from pmCherry-N1 (Clontech) with PW_GG_SV40ori_F and PW_GG_SV40ori_R and cloned in pGGF000 to create pPW53.

eGFP coding sequence was amplified from pGGD001 with oGD261 and oGD262 and cloned in pGGD000 to create pGD165. Clover coding sequence was amplified from Clover-mRuby2-FRET-10 (M. Davidson, Addgene) with oGD317 and oGD318 and cloned in pGGD000 with a linker made up by annealing oGD315 and oGD316 (D-TGCA linker) to create pGD250. mCerulean3 coding sequence was amplified from pmCer3-N1 (Markwardt et al., 2011) with oGD317 and oGD318 and cloned in pGGD000 with the D-TGCA linker to create pGD251. T-Sapphire coding sequence was amplified from PstI-digested p35S::TSapphire-mOrange-nos plasmid (Bayle et al., 2008) with oGD317 and oGD318 and cloned in pGGD000 with the D-TGCA linker to create pGD252. mTurquoise2 was amplified from mTurquoise2-N1 (Goedhart et al., 2012) and cloned in pGGD000 with the D-TGCA linker to create pGD425. mRuby2 coding sequence was amplified from Clover-mRuby2-FRET-10 with oGD325 and oGD326 and cloned in pGGD000 to create pGD253. mRuby3 was amplified from pNCS-mRuby3 (Bajar et al., 2016a) with oGD325 and oGD318 and cloned into pGGD000 to create pGD341. mOrange coding sequence was amplified from p35S::TSapphire-mOrange-nos with oGD318 and oGD331, followed by a second PCR with oGD327 and oGD318 and cloned in pGGD000 to create pGD254. Venus coding sequence was amplified from pABindVenus (Bleckmann et al., 2010) with oGD317 and oGD318 and cloned in pGGD000 with the D-TGCA linker to create pGD255. mNeonGreen coding sequence was amplified from pNCS-mNeonGreen (Allele Biotechenology) with oGD343 and oGD344 and cloned in pGGD000 with the D-TGCA linker to create pGD352. mCherry coding sequence was amplified from pGGC015 with RD_GG_mCherry_C-tag_F and RD_GG_mCherry_C-tag_R cloned in pGGD000 to create pRD53. mScarlet coding sequence was amplified from pmScarlet-C1 (Bindels et al., 2017) with RD_GG_mScarlet_C-tag_R and RD_GG_mScarlet_C-tag_F cloned in pGGD000 to create pRD134. mKate2 coding sequence was amplified from pmKate2-C1 (Shcherbo et al., 2009) with RD_mKate2_GG_C-tag_R and RD_mKate2_GG_C-tag_F cloned in pGGD000 to create pRD141.

### Generation of Transient Expression Plasmids

The plasmid backbone containing the XVE expression cassette under the control of the 35S promoter and RBCS terminator, and the A and G GreenGate cloning sites (pGD283) was generated by combining the inserts from pGGA004 (p35S), pGGB002 (Omega element), pRD71 (XVE), pGGD002 (N-dummy), pGGE001 (tRBCS), pGGG001 (F-H adapter), and a double-stranded methylated linker containing H and G GreenGate sites (Supplementary table 1) into pGGZ001 in a single step GreenGate reaction (Lampropoulos et al., 2013).

Individual expression cassettes were prepared as intermediate plasmids. Donor constructs were prepared by combining pBLAA001 (pLexA-mini35S), pGGB002, pGD293 (CRNΔKi), fluorophore entry plasmid, pGGE009 (tUBQ10), and pGGG001 (F-H adapter) into pGGM000 in a single step GreenGate reaction. For the acceptor constructs, pGD293 was replaced by pGD292 (CLV2) or pGD318 (myr) and pGGG001 by pGGG002 (H-A adapter), pPW53 was added and pGGN000 was used as a destination plasmid. For the donor and acceptor only constructs, untagged CRNΔKi and CLV2 expression cassettes were generated using pGGD002 in place of a fluorophore entry plasmid.

Transient expression plasmids containing both donor and acceptor fusion proteins were generated by combining the donor and acceptor intermediate plasmids in the pGD283 backbone. For the expression of a single fusion protein either the donor or acceptor intermediate plasmids were replaced with linkers with A-H or H-G overhangs (Supplementary table 1), respectively.

### Transient Expression in Nicotiana

*Nicotiana benthamiana* infiltration was carried out using standard protocole (Li, 2011). Briefly, agrobacterium strain C58:pmp90:pSOUP carrying the expression plasmid were cultivated overnight in dYt medium. Cells were then pelleted and resuspended in infiltration medium (MgCl_2_ 10mM; MES-K 10mM pH5.6; 150μM acetosyringone) to an OD_600_ of 0.4 and incubated for 2h at room temperature. Bacteria solutions were then mixed in equal quantities with Agrobacterium strain GV3101:p19 expressing the p19 silencing inhibitor. For the co-infiltration of CRNΔKi-eGFP and CLV2-mCherry or CLV2 untagged, bacteria were resuspended in infiltration medium to an OD_600_ of 0.6 and mixed in 1:3 ratio with GV3101:p19 cells. Bacteria mixes were then infiltrated to the abaxial side of 3-4 weeks old *Nicotiana benthamiana*. After 2-4 days under continuous light, protein expression was induced overnight by spraying the abaxial side of infiltrated leaves with an estradiol solution (Estradiol 20μM; Tween-20 0.1%).

### Fluorescence Imaging

Fluorescence imaging was performed with a Zeiss LSM 780 confocal microscope (40x Water immersion objective, Zeiss C-PlanApo, NA 1.2). T-Sapphire was excited at 405 nm; mCerulean3 and mTurquoise2 at 458 nm; Clover, mNeonGreen, and eGFP at 488 nm; Venus at 514nm, and mCherry, mRuby2, mRuby3, mScarlet, mKate2 and mOrange at 561 nm. Signal for each fluorophore was recorded within the maximum emission peak while avoiding auto-fluorescence above 650nm. The fluorescent properties of the fluorophores used here are available as “protein collection” on the website fpbase.org (https://www.fpbase.org/collection/332/).

### Time-Correlated Single Photon Counting

Fluorescence lifetime was acquired with a Zeiss LSM 780 confocal microscope (40x Water immersion objective, Zeiss C-PlanApo, NA 1.2). Time correlated single photon counting was performed with picosecond resolution (PicoQuant Hydra Harp 400). Fluorophores were excited with either a 440nm (LDH-D-C-440, 32 MHz) or 485 nm (LDH-D-C-485, 32 MHz) pulsed polarized diode laser with a power of 1 μW at the objective lens. Emitted light was separated by a polarizing beam splitter and parallel and perpendicular photons were selected with a fluorophore-specific band-pass filter (Supplementary table 2) and detected with Tau-SPADs (PicoQuant) simultaneously for the acceptor and donor channels. Image acquisition was done at zoom 8 with a resolution of 256×256 pixel with a pixel size of 0.1μm a dwell time of 12.6 μs, and photons were collected over 60 frames. Special care was taken during imaging to avoid chloroplasts-containing regions and cells with very high donor expression to avoid pile-up effect.

### Fluorescence Decay Analysis

Fluorescence decay was analysed in Symphotime 64 (version 2.4; PicoQuant) using the Lifetime FRET Image analysis tool. Only data from the donor parallel channel were kept for the analysis. TCSPC channels were binned by eight, count threshold was set so that the background was removed, and chloroplasts were manually removed. Additionally, in case some pixels were above the pile-up limit (10% of the laser repetition rate, *i.e*. 2421 counts), they were manually removed; counts values were in most cases below 5% of the laser repetition rate. Internal response function was determined by measuring the fluorescence decay of saturated erythrosine, or Atto425 dye for blue donors, quenched in saturated KI using the same hardware settings as for the FRET pair of interest. Fluorescence decay was fitted using a multi-exponential decay and the amplitude-weighted lifetime was considered as the sample’s apparent lifetime. FRET efficiency was calculated as the lifetime of the FRET sample over the arithmetic mean of the lifetimes of the donor only samples measured in the same experiment: 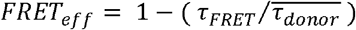. All measurements were done in at least two independent experiments.

### Fluorescence Intensity Measurement

To calculate the A/D ratio, donor and acceptor were excited with a 485 nm pulsed laser (LDH-D-C-485, 32 MHz, 1μW) and intensities were recorded with Tau-SPADs (PicoQuant) in the setup described above. Fluorescence intensity was extracted using FIJI (Schindelin et al., 2012; Rueden et al., 2017). Membrane regions were selected by thresholding the donor channel and the same ROI was applied on the acceptor channel and the signal’s integrated density was measured in the ROI.

Background fluorescence was similarly measured by imaging mock-infiltrated plants and extracting the signal intensity over the whole image in the donor channel. Likewise, bleed-through measurements were performed on plants co-expressing CRNΔKi-untagged and CLV2-donor constructs. Donor intensities were calculated in the same way using donor only samples.

For photobleaching, plants co-expressing either CRNΔKi-donor and CLV2-untagged or CRNΔKi-untagged and CLV2-donor were excited with the donor excitation laser and fluorescence was recorded as for a lifetime measurement. A ROI was set over the membrane and fluorescence intensity was recorded in every frame.

## Authors contributions

GD conceived the project and designed the experiments with input from all authors. GD prepared the figures and wrote the manuscript with the help of RS. GD and PS created expression vectors and performed measurements. GD, PS, and SH analyzed the data. SH and SWP provided critical help with lifetime measurement and analysis. All authors discussed the results and commented on the manuscript.

## Supplementary material

Supplementary figure 1. Myristoylated-mCherry co-localizes with CRNΔKi-eGFP.

Supplementary figure 2. Lifetime stability for each donor.

Supplementary table 1. Oligonucleotides used in this study.

Supplementary table 2. Microscope configuration for each FRET pair.

Supplementary table 3. Plasmids available from Addgene.

## Material availability

Plasmids listed in Supplementary table 3 are available from Addgene (XXXX). All other materials are available upon request.

## Acknowledgments

The work of all authors was supported by the DFG through CRC1208. We wish to thank R. Burkart and A. Bleckmann for sharing unpublished plasmids, and Y. Stahl and M. Somssich for useful discussion. Clover-mRuby2-FRET-10 was a gift from Michael Davidson (Addgene plasmid # 58169; http://n2t.net/addgene:58169; RRID: Addgene_58169).

