## Supplementary information for "Over the rainbow: a practical guide for fluorescent protein selection in plant FRET experiments"

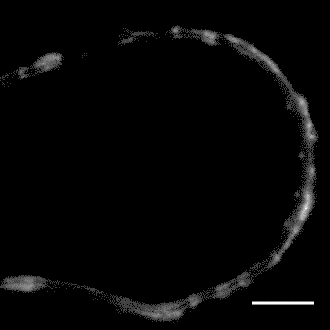

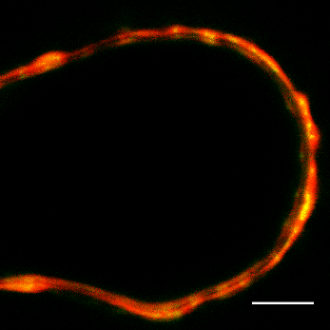

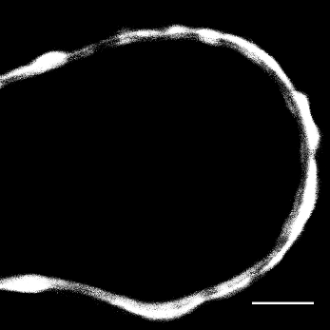

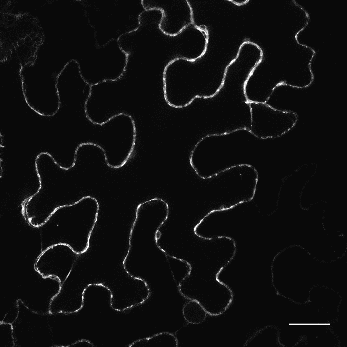

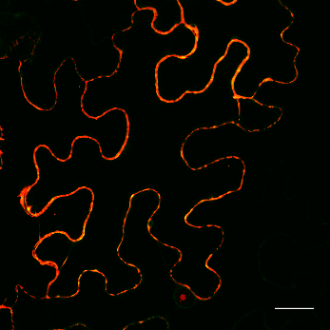

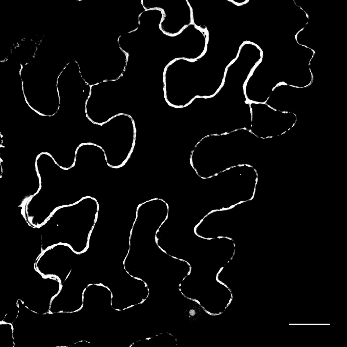

CRNΔKi-eGFP

myr-mCherry

Merge

**A**

**B**

**Supplementary figure 1. Myristoylated-mCherry co-localizes with CRNΔKi-eGFP.** Confocal microscopy images of Benthamiana epidermis cells co-expressing CRNΔKi-eGFP (left), CLV2-untagged, and myr-mCherry (middle) constructs. False color merged image (right) shows co-localization of both eGFP and mCherry at the plasma-membrane (green: eGFP; red: mCherry) in whole cells (A) and in close-up of a membrane section (B). Scale bar: 25µm (top), 5µm (bottom) (C) Acceptor/Donor intensity ratios in the cells used for fluorescence lifetime measurement.

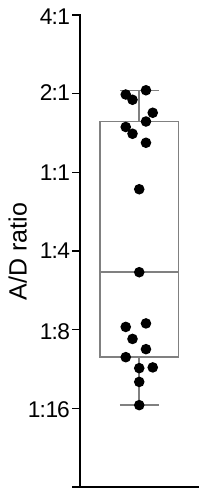

**C**

**Supplementary figure 2. Lifetime stability for each donor.**
Distance of the donor only samples from the population means. Each point represents an individual sample. Bars represent the mean distances.

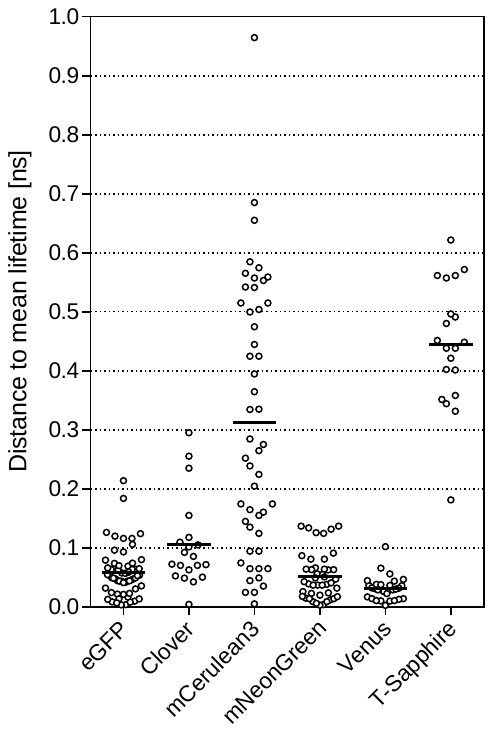

**Supplementary table 1.** Oligonucleotides used in this study. [mC] indicates the presence of a 5-methyl-dCytosine.

| Name | Target | Sequence |
| --- | --- | --- |
| GG_XVE_F | XVE | AAAGGTCTCAGGCTATGAAAGCG |
| G_XVE_R | XVE | AAACACTGAGACTGTGGCAGGG |
| PW_XVE_3084_AT_R | XVE | GGAGCGCCAGACGAGTCCAATCATCAGGAT |
| PW_XVE_3084_TA_F | XVE | ATCCTGATGATTGGACTCGTCTGGCGCTCC |
| GG_EST_F | LexA-mini35S | AACAGGTCTCAACCTTGCATGCCAGCTTGGGCTGCAGGTCGAGGCT |
| GG_EST_R | LexA-mini35S | AACAGGTCTCTTGTTCTTCAGCGTGTCCTCTCCAAATGAAATGAA |
| PS-GG-CDS-Clv2-F | CLV2 | AAAGGTCTCAGGCTTAATGATAAAGATTGCAG |
| PS-GG-CDS-Clv2-R | CLV2 | TTTGGTCTCACTGAGAAAGCTGGGTAAG |
| oGD335 | CRNΔKi | AAAGGTCTCAGGCTTAATGAAGCAAAGAAGAAGAAG |
| oGD336 | CRNΔKi | AAAGGTCTCACTGATAACACCAAAGCTGAAACC |
| oGD339 | myr | GGCTCTATGGGAAGAAGAAAAAGAAAACCTAAA |
| oGD340 | myr | CTGATTTAGGTTTTCTTTTTCTTCTTCCCATAG |
| PW_GG_SV40ori_F | SV40ori | AAAGGTCTCAACTAGGTGTGGAAAGTCCCC |
| PW_GG_SV40ori_R | SV40ori | AAAGGTCTCAATACGGCCTCCAAAAAAGCC |
| oGD261 | eGFP | AAAGGTCTCATCAGGCAGCGGCTCTGGATCG |
| oGD262 | eGFP | ATAGGGCGAGAATTCGGTCTCAGC |
| oGD317 | Clover,  mCerulean3,  Venus,  T-Sapphire,  mTurquoise2 | AAAGGTCTCATGCAATGGTGAGCAAGGGCG |
| oGD318 | Clover,  mCerulean3,  Venus,  mOrange,  T-Sapphire,  mTurquoise2, mRuby3 | AAAGGTCTCAGCAGTTACTTGTACAGCTCGTCCATGC |
| oGD315 | D-TGCA linker | TCAGGCAGCGGCTCTGGATCGGCGGCCGC |
| oGD316 | D-TGCA linker | TGCAGCGGCCGCCGATCCAGAGCCGCTGCC |
| oGD325 | mRuby2, mRuby3 | AAAGGTCTCATCAGGAATGGTGTCTAAGGGCGAAGAG |
| oGD326 | mRuby2 | AAAGGTCTCAGCAGTTACTTGTACAGCTCGTCCATCCC |
| oGD331 | mOrange | TTGTACAAAGTGGTTGATGGG |
| oGD327 | mOrange | AAAGGTCTCATCAGGAATGGTGAGCAAGGGCGAG |
| oGD343 | mNeonGreen | AAAGGTCTCATGCAATGGTGAGCAAGGGAGAG |
| oGD344 | mNeonGreen | AAAGGTCTCAGCAGTTACTTGTAAAGCTCGTCCATTC |
| RD_GG_mCherry_C-tag_F | mCherry | AAAGGTCTCATCAGCAATGGTGAGCAAGG |
| RD_GG_mCherry_C-tag_R | mCherry | AAAGGTCTCAGCAGTTACTTGTACAGCTCGTC |
| RD_GG_mScarlet_C-tag_R | mScarlet | AAAGGTCTCAGCAGTTACTTGTACAGCTC |
| RD_GG_mScarlet_C-tag_F | mScarlet | AAAGGTCTCATCAGTTATGGTGAGCAAG |
| RD_mKate2_GG_C-tag_R | mKate2 | AAAGGTCTCAGCAGTTAGCGGTGAC |
| RD_mKate2_GG_C-tag_F | mKate2 | AAAGGTCTCATCAGTTATGGTGTCGG |
| linker_AH_met_F | A-H methylated linker | ACCTACCTTGAGAC[mC]GAAAAGGTGGTCT[mC]A |
| linker_AH_met_R | A-H methylated linker | CCTATGAGAC[mC]ACCTTTTCGGTCT[mC]AAGGT |
| linker_HG_met_F | HG methylated linker | TAGGACCTTGAGAC[mC]GAAAAGGTGGTCT[mC]A |
| linker_HG_met_R | HG methylated linker | ATACTGAGAC[mC]ACCTTTTCGGTCT[mC]AAGGT |

**Supplementary table 2.** Microscope configuration for each FRET pairs.

| Donor | Acceptor | Donor Ex/Em peaks | Acceptor Ex/Em peaks | Excitation laser [nm] | Donor band-pass filter | Acceptor band-pass filter | Beamsplitter |
| --- | --- | --- | --- | --- | --- | --- | --- |
| eGFP | mCherry | 488/507 | 587/610 | 485 | 520/35 | 607/70 | LP560 |
| eGFP | mScarlet | 488/507 | 569/594 | 485 | 520/35 | 607/70 | LP560 |
| Clover | mRuby2 | 505/515 | 559/600 | 485 | 520/35 | 607/70 | LP560 |
| mNeonGreen | mRuby2 | 506/517 | 559/600 | 485 | 520/35 | 607/70 | LP560 |
| T-Sapphire | mOrange | 399/511 | 548/562 | 440 | 520/35 | 607/70 | LP560 |
| Venus | mKate2 | 515/527 | 588/633 | 485 | 534/30 | 607/70 | LP560 |
| mCerulean3 | Venus | 433/475 | 515/527 | 440 | 482/35 | 534/30 | LP510 |
| mCerulean3 | mNeonGreen | 433/475 | 506/517 | 440 | 482/35 | 520/35 | LP510 |
| mTurquoise2 | Venus | 434/474 | 515/527 | 440 | 482/35 | 534/30 | LP510 |
| mTurquoise2 | mNeonGreen | 434/474 | 506/517 | 440 | 482/35 | 520/35 | LP510 |
| mNeonGreen | mRuby3 | 506/517 | 558/592 | 485 | 520/35 | 607/70 | LP560 |
| Venus | mRuby3 | 515/527 | 558/592 | 485 | 520/35 | 607/70 | LP560 |

**Supplementary table 3.** Plasmids available from Addgene. All plasmids are suitable for use with the GreenGate kit (Lampropoulos et al., 2013).

| Content | Reference | Backbone |
| --- | --- | --- |
| *Destination plasmid* | | |
| 35S:XVE:tRBCS | pGD283 | pGGZ001 |
| *Entry plasmids* | | |
| pLexA-mini35S | pBLAA001 | pGGA000 |
| myr | pGD318 | pGGC000 |
| eGFP | pGD165 | pGGD000 |
| Clover | pGD250 | pGGD000 |
| mCerulean3 | pGD251 | pGGD000 |
| T-Sapphire | pGD252 | pGGD000 |
| mRuby2 | pGD253 | pGGD000 |
| mOrange | pGD254 | pGGD000 |
| Venus | pGD255 | pGGD000 |
| mCherry | pRD53 | pGGD000 |
| mScarlet | pRD134 | pGGD000 |
| mKate2 | pRD141 | pGGD000 |
| mNeonGreen | pGD352 | pGGD000 |
| mTurquoise2 | pGD425 | pGGD000 |
| mRuby3 | pGD431 | pGGD000 |
